# Towards automatic derivation of geometry-based descriptors as surrogates for complex computational approaches in enzyme-substrate prediction

**DOI:** 10.1101/2025.11.26.690723

**Authors:** Carlos Sequeiros-Borja, Petr Škoda, Jan Brezovsky

## Abstract

Accurate prediction of enzyme-substrate interactions remains a fundamental challenge in biocatalysis and drug discovery. While machine learning approaches have shown promise, they require extensive training data and often lack mechanistic interpretability. Here, we present a novel methodology that automatically derives geometry-based descriptors from enzyme-substrate complex structures to predict substrate specificity. Our approach simplifies complex catalytic mechanisms into interpretable geometric filters comprising critical inter-atomic distances and accessibility of atomic pairs parameters. We validated this methodology using two mechanistically distinct enzyme families with minimal training data: haloalkane dehalogenases (9 enzymes and 53 substrates) and aldehyde reductases (9 enzymes and 36 substrates). The filters demonstrated robust performance across chemically diverse substrates. On testing datasets, the derived filters achieved average accuracy of 77% and sensitivity of 94% for haloalkane dehalogenases and average 57% recall of true substrates for aldehyde reductases, exceeding state-of-the-art machine learning methods for substrate predictions on these datasets. Crucially, the geometric descriptors directly correspond to catalytic requirements, providing mechanistic insights into substrate recognition. This interpretable, mechanism-based approach requires minimal training data and can be readily applied to newly characterized enzymes, offering a powerful tool for enzyme engineering and substrate screening applications.

## Introduction

Enzymes are natural and efficiently evolved catalysts able to perform a myriad of chemical reactions such as the hydrolysis of substrates,^[1]^ removal and transfer of chemical groups,^[2]^ enantioselectivity,^[3]^ cleavage of bonds and isomerization among others.^[4]^ Given the plasticity and usefulness of enzymes to perform desired chemical reactions, these molecules have been, and still are, intensively studied in different fields from industry or science, like biotechnology, bioengineering, evolution and biochemistry.^[5–7]^ In this sense, the elucidation of the reaction mechanism of enzymes is essential, since the information on their reaction can be used to obtain desired end products or subproducts. This can be performed with experimental, theoretical (computational) or a combination of both methodologies.^[8–10]^

While experimental approaches remain valuable, computational methods have become vital for understanding enzymatic mechanisms. Among the computationally most common ones are Empirical Valence Bond^[11,12]^ and hybrid quantum mechanical (QM)/molecular mechanical^[13,14]^ approaches. In the last decades, the prediction of substrates for known enzymes has attracted a lot interests with the increased use of Machine Learning (ML) techniques and more recently with Artificial Intelligence (AI.^[15–19]^ In the area of computational prediction of substrates, we can identify three types: sequence-based, structure-based, and a combination of both.^[18–22]^ Lately, sequence-based tools have taken advantage of the vast amount of sequence data present in public databases, and huge advances of ML and AI techniques. For instance, the ESP method converts the amino acid sequence into a numerical representation through a modified ESM-1b transformer model, and the candidate substrates into fingerprints derived from a graph neural network.^[19]^ Similarly, FusionESP uses two different encoders to represent the enzyme and substrate independently to high-dimensional vectors, which are later transformed in combinations to yield a numerical value between 0 and 1 to determine a possible interaction.^[18]^

In contrast to sequence-based approaches, methods that use structural information as main source of information frequently employ expensive computational calculations and rely on a lower amount of available data, usually in form of 3D structures of complexes, compared to sequence-based ones. As the name suggests, structure-based methodologies require 3D structures of the enzymes and substrates to operate, and such information can be obtained through experimental or computational methods. Experimental 3D structures of enzymes bound to their substrates are arguably the most reliable source of structural information, however, this approach has considerable limitation due to high costs of materials, equipment and time required to obtain a single structure of ES-complex. Hence, molecular docking is mostly used as an attractive approach to obtain acceptable structures of ES-complexes at a relatively low cost.^[23–25]^ Alternatively, the 3D structures of complexes could be generated with co-folding AI models like, AlphaFold3,^[26]^ RoseTTAFold All-Atom,^[27]^ Chai-2,^[28]^ and Boltz-2.^[29]^ The utilization of these models have clear benefit of full structure flexibility, but might lead to generation of highly unphysical structures especially when operating on complexes outside their training data.^[30]^ One of the structure-based methodology that has been proved to be successful for ligand prediction is Quantitative Structure Activity Relationship (QSAR), which builds a model in several steps, initially gathering chemogenomic data from different databases, then several chemical descriptors are determined at different levels of representation depending on the approach employed (ranging from 1D to nD), to finally correlate these descriptors to a defined biological property using ML techniques.^[31,32]^ Although this methodology and others produce a deep understanding about the reaction mechanism at atomic level, and provide models for prediction of inhibitors and substrates, it requires a 3D structure of the enzyme-substrate (ES) complex, quite substantial computational resources, high-level knowledge on the reaction system and expertise in structural bioinformatics. Another alternative is to analyze ES-complexes with various geometric descriptors that were developed to capture essence of the reaction geometry to flag a complex as potentially reactive or not. Previous work has shown that the use of geometric descriptors can be a fruitful method for estimating substrate’s reactivity for various enzyme classes.^[33–35]^ The main drawback of these studies is that the geometric descriptors were derived a priori based on extensive QM calculations on a given enzyme family, limiting their widespread adoption. This creates a significant barrier: while geometric approaches offer mechanistic interpretability and require less training data than most ML methods, their development has remained prohibitively expensive and specialized, preventing application to the vast majority of enzyme families.

To address this gap, and building on structures of ES-complexes, we developed a methodology capable of simplifying the structural characteristics of an enzyme’s reaction mechanism into a minimal subset of relevant geometric descriptors (**Figure 1**), employing the knowledge available in literature, public databases and ML techniques. In our approach, we obtain such geometric descriptors based on the automated analysis of recurring binding patterns of known substrates across different family members, avoiding the difficult task of mechanistic studies of the reaction. Hence, our approach does not require specific knowledge about the reaction mechanism and harnesses unconstrained molecular docking as source of the structures of ES-complexes. The only necessary input is the definition of the residues that are expected to interact directly with substrates. Initially, we developed and tested our approach for the haloalkane dehalogenase (HLD) family (EC 3.8.1.5), since it comes with a considerable amount of structural data, and geometric approaches have previously been described for this family.^[36–39]^ To further test the generalization of the method and its limits, we also used the aldehyde reductase (AldR) NADPH dependent family (EC 1.1.1.2), since the presence of a cofactor for the reaction markedly increases the complexity.

**Figure 1.**
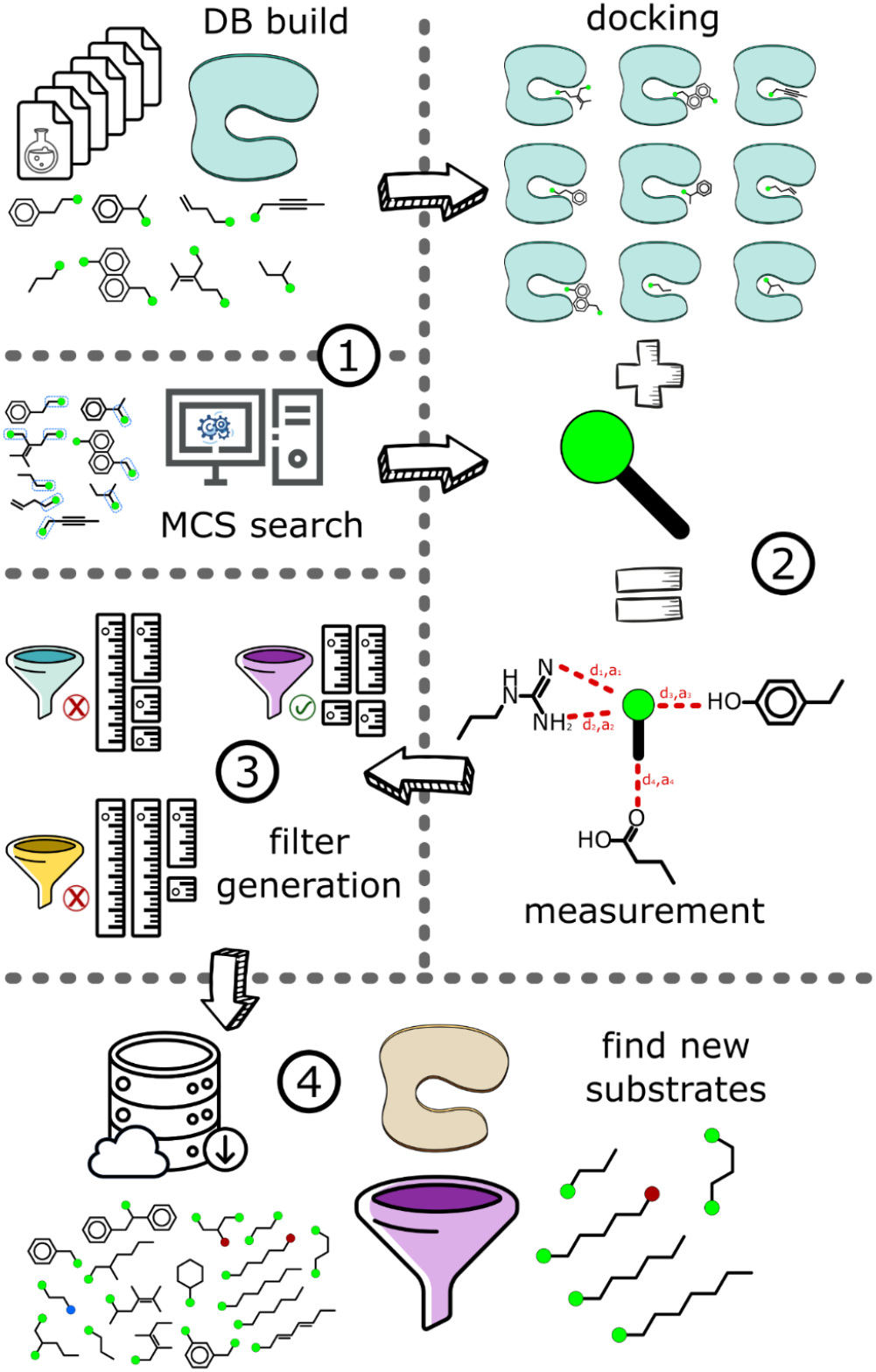
The substrate prediction workflow based on family-wide geometry descriptors. The whole process can be summarized in four stages. 1) Enzyme-substrate (ES) pairs are gathered from databases and the literature. With this information, the structures of substrates are clustered and their maximum common substructure (MCS) is identified as their reactive fragment. 2) Structures of these ES-complexes are prepared via molecular docking calculations and the arrangements between atoms of reactive fragment and the protein active site residues (distances and accessibility) are measured for each ES-complex conformation. 3) The ES-complex conformations are clustered using the measurements as input. From these clusters, the geometry descriptors are derived, to be used as a filter in substrate predictions. 4) New potential substrates that contain the reactive fragment are obtained from public databases and docked into a target enzyme. The filter selected in the previous stage is used to discriminate between suitable substrates and inactive compound for the target enzyme.

### Computational Methods

Enzyme-substrate pairs were compiled from BRENDA database ^[40]^ and literature for HLDs (**Supporting Information Table 1**) and AldRs (**Supporting Information Table 2**). Protein structures were obtained from PDB^[41,42]^ or AlphaFold database^[43,44]^ when experimental structures were unavailable; substrate structures were retrieved from PubChem^[45]^ (**Supporting Information Tables 3 and 4**). DmmA (PDB: 3U1T)^[46]^ and AKR1B14 (PDB: 3O3R)^[47]^ enzymes served as test cases for HLDs and AldRs, respectively. To determine the predictive performance of the method, it would be ideal to have both true positives and true negatives, which are unfortunately not available for every family. In the case of 3U1T, there we compiled 8 true negatives and 13 true positives, while for 3O3R, there were only 25 true positives (**Supporting Information Table 5**). Given such dataset compositions, the accuracy was calculated for HLD, while the recall was obtained for AldR. For the evaluation of sensitivity of the filter performance to the composition of input substrates, the available substrates were divided into five substrate groups (SG), and the filters were generated while leaving-one-SG-out. The separation was performed with AgglomerativeClustering method of SciKitLearn package, with the default settings except the compute distances parameter set to true. To create the matrix for the clustering, the Daylight-like fingerprint of each substrate was obtained with the RDKit package, and the Tanimoto similarity was calculated for each combination of two substrates. For substrate clustering, also the candidate substrates of the target enzyme were included but were not used in the filter generation.

Protein structures were prepared by removing water molecules and non-protein atoms (except NADPH cofactor for AldRs), selecting the highest-occupancy conformer for alternate positions, and aligning all structures within each family to their respective target enzymes (3U1T or 3O3R). Then, protein and substrate structures were prepared for docking with MGLTools v1.5.7 package.^[48]^ For the proteins, the prepare_receptor4.py script with flags ‘checkhydrogens’ and ‘nphs_lps’ active was employed, while for substrates, the prepare_ligand4.py script with default settings was used. For each enzyme family, an unified docking box center on active site residues retrieved from M-CSA database^[49]^ was used. The box size was adjusted to cover entire set of active sites. Molecular docking was performed with Autodock Vina v1.1.2,^[50]^ which was modified to generate an increased number of binding poses, enabling identification of binding modes shared among all substrates. Additionally, to avoid any artifact produced by randomization within the docking software, the docking experiments for all ES-complexes were performed only once, and these resulting conformations were reused throughout the analysis.

The developed approach assumes that all substrates recognized by the explored enzyme family share one chemical substructure that is entering the catalytic reaction and this substructure interacts with catalytic groups of enzymes via conserved binding geometry. To identify the relevant fragments in substrate molecules, we searched for maximum common substructure (MCS) among the substrates of a given enzyme family using the RDKit v2023.03.1 package.^[51]^ The FindMCS function of this package was modified to recognize halogen atoms as a single group and to match the valences of the atoms. In some substrates, multiple MCS were identified. So, for each substrate docked to an enzyme, we have multiple conformations/poses and in each of the poses, we recognize at least one fragment defined by MCS.

Next, all docked fragment poses were filtered to retain only those with energetically favorable Vina scores, fragment atoms within 8 Å of the closest catalytic residues, and appropriate electrostatic complementarity of fragment atoms with the selected functional groups of catalytic residues. For retained fragments, distances between all fragment atoms and functional groups of catalytic residues (defined by M-CSA, Supporting information table 6) were calculated for each docked conformation, creating a uniform distance descriptor vector containing measured distances for all retained interaction pairs.

Fragment poses were clustered by their distance profiles using a two-stage approach with the SciKitLearn v1.1.2^[52]^ and hdbscan v0.8.40 packages.^[53]^ Here, HDBSCAN (minimum cluster size=10, ε=1.0) was applied to determine cluster number, followed by KMeans for final assignment to prevent reactive-like poses (often low-population) from being discarded as noise. Each produced cluster contains substrate fragments that share similar binding geometries and interaction patterns with enzyme active sites. Inside each cluster, several fragment conformations from the same ES-pair could be present. Hence, for each cluster, representative poses were selected using a simplified Coulomb score that captures the electrostatic complementarity between the substrate’s fragment and functional groups of enzyme active site, providing a relative measure of electrostatic favorability across different binding poses. To finalize the set of available pairwise descriptors for each fragment, atom pairs with average distances >6 Å were excluded and, for retained pairs, accessibility was calculated by ray-tracing from enzyme atoms to fragment atoms, quantifying the fraction of substrate atom surface accessible to each catalytic group.

In the last step, geometry-based filters of enzyme-substrate activity were generated through statistical analysis of successful binding modes, implementing an iterative relaxation algorithm to define permissible ranges for key molecular interactions. For each cluster of fragment poses, two categories of filters were derived simultaneously: distance-based filters defining acceptable ranges for atom-atom distances between substrate fragments and catalytic residues, ii) and accessibility filters ensuring sufficient exposure of substrate atoms to reactive centers. The algorithm employs a greedy optimization strategy, utilizing normalized error functions to balance contributions from both filter types, with the normalization scheme ensuring comparable weight across distance and accessibility metrics. Initially, the limits were at most 0.0 Å and at least 100 % for the distance and accessibility, respectively. At each iteration, the fragment pose exhibiting the minimum total squared error across both filters is selected for inclusion. This fragment represents the binding pose closest to satisfying all geometric and accessibility constraints without modification. The specific filter preventing this fragment’s acceptance (identified by its minimum individual error contribution) is then relaxed by adjusting its interval boundaries to exactly accommodate the fragment’s value. This process ensures that filters are minimally modified to achieve the target coverage of 50% of input fragments, meaning that such geometric criteria for reactivity are the most stringent yet fulfilled by majority of ES-complexes present in each cluster of binding poses, i.e. common binding pose of enzyme family. When more than one filter was generated per cluster, the number of ES-fragments used for its generation was used as criterion to select the more general filter.

After all the filters were generated, the candidate substrates followed the same process of docking and the measures for atom pairs were taken in the same way as for the initial substrates, with the only difference that three independent rounds of docking were performed to increase the available poses. The candidate substrates with any conformation fulfilling both filter criteria were considered as likely substrates of target enzyme.

The python implementation of the filter derivation and candidate screening, all generated dataset on HLD and AldR families are available from Zenodo repository (https://zenodo.org/records/10901578) to support reproducibility and reusability of both the data and method.

## Results and Discussion

The HLD family was selected as the primary test case for our methodology due to its well-characterized substrate specificity and extensive experimental validation. HLDs exhibit broad substrate specificity while maintaining a conserved catalytic mechanism,^[54,55]^ which coupled with availability of multiple high-quality crystal structures makes them fitting candidates for developing generalizable geometry-based descriptors. Additionally, the industrial relevance of HLDs in bioremediation and biosynthesis applications underscores the practical importance of accurate substrate prediction for this enzyme family.^[56,57]^ Importantly, this enzyme family has been previously explored using geometric criteria derived based on QM calculations,^[33]^ providing a robust benchmark for the comparison of current approach.

The training dataset for the HLD family comprised 9 enzymes with experimentally verified substrates obtained from BRENDA and literature sources (**Supporting Information Table 1**). A total of 74 unique substrates were categorized into five substrate groups (SG0-SG4) based on their chemical structures, with 53 substrates designated for training and 21 for testing. The molecular docking procedure generated between 14,000-25,000 ES-poses, which were subsequently clustered into 2-5 distinct binding modes based on their interaction distance profiles. From these binding modes, mostly a single geometric filter was derived for each substrate group exclusion experiment, with the best-performing filter selected based on the number of unique ES-complexes used for its derivation (**Supporting Information Table 7**). To evaluate the robustness of filter performance irrespective of substrate diversity, we employed a leave-one-SG-out strategy. Each of the five SGs was systematically excluded from the training set, and filters were generated using the remaining four SGs (**Table 1 and Supporting Information Table 1)**. The derived filters demonstrated remarkable consistency across different training datasets, with geometric descriptors primarily focusing on nucleophilic aspartate, and halide-stabilizing asparagine and tryptophan (**Figure 2**). The interacting atoms are required to have a high accessibility (∼0.8 and more) and the distances below ∼ 3.7 Å in the most lenient case (**Supporting Information Table 8**). This consistency in filter composition across different training sets indicates the robustness of our approach in identifying mechanistically relevant geometric constraints.

**Figure 2.**
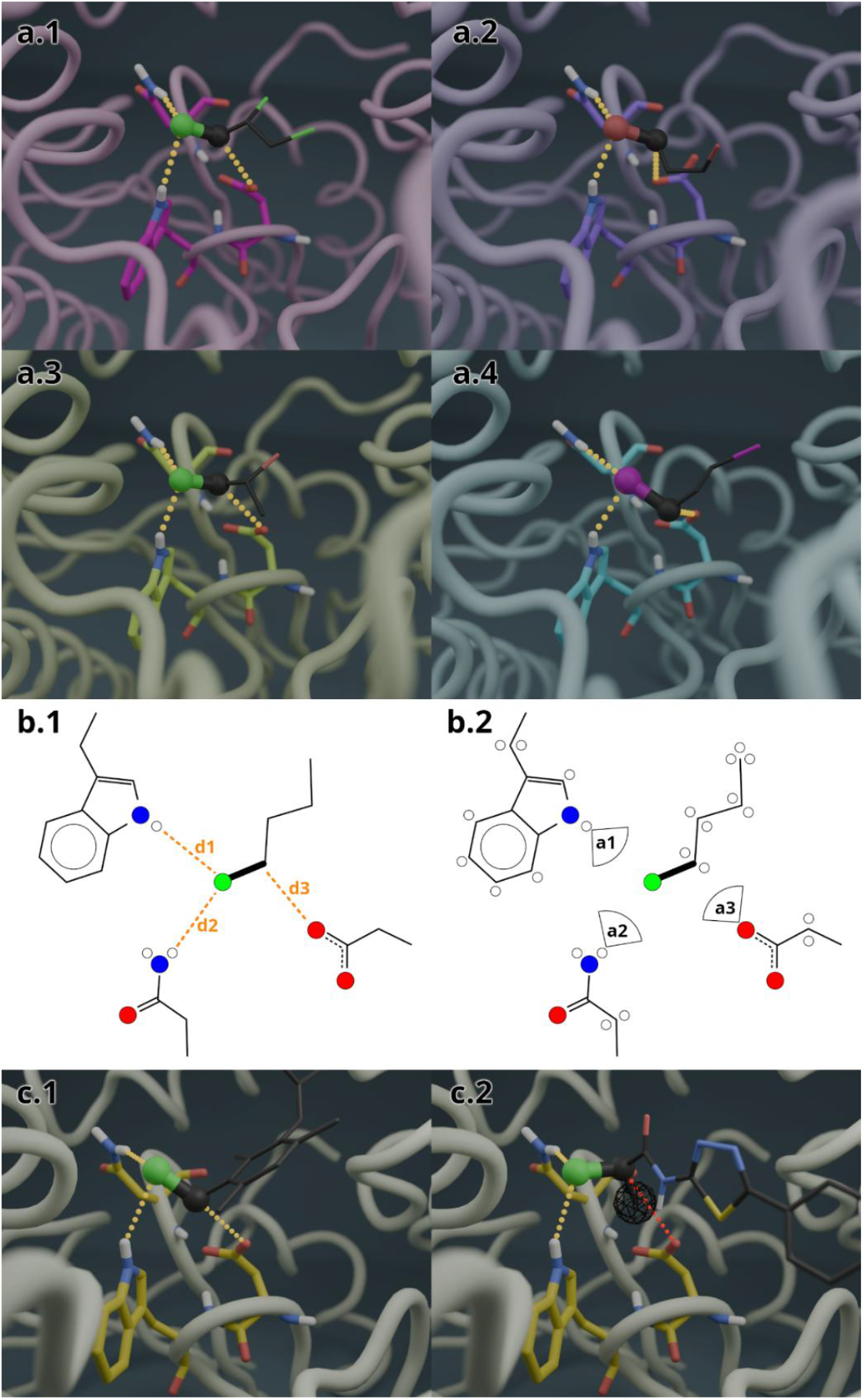
Geometric descriptors for HLD substrate recognition. a) Representative ES-complexes of the HLD family with verified substrates showcasing the geometric descriptors. ES-complexes as PDB and CID IDs are presented in order 1CQW–7285 (a.1), 1MJ5–8001 (a.2), 2QVB– 18175 (a.3), and 3A2M–12314 (a.4). Proteins represented as ribbon with the key catalytic residues ASN, ASP and TRP as sticks. Substrates in lines representation with the reactive fragment atoms as ball–stick, and the geometric descriptors as dotted yellow lines. b) Schematic representation of the geometric filter criteria derived from training data. Key atom pairs are shown with their distance cons traints (b.1) and accessibility requirements (b.2). Atom pairs include ASN-78 HD21/HD22 to halogen, ASP-144 OD1/OD2 to carbon, and TRP-145 HE1 to halogen, numbered according to 3U1T structure. c) Side-by-side comparison of a correct binding pose (c.1) that satisfies all geometric filters (distances within range, high accessibility) versus an incorrect binding pose (c.2) that fails the accessibility criterion for the ASP-144 OD1/OD2 to carbon interaction in the target enzyme 3U1T. Protein in ribbon representation with the key residues as sticks. Ca ndidates shown as lines with the reactive fragment atoms as ball–stick, and the geometric descriptors as dotted yellow lines. In c.2, carbon atom of the candidate shielding the accessibility represented as wired sphere, with its corresponding hindered interaction as dotted red line. This illustrates how geometric filters discriminate between productive and non-productive binding modes.

**Table 1.**
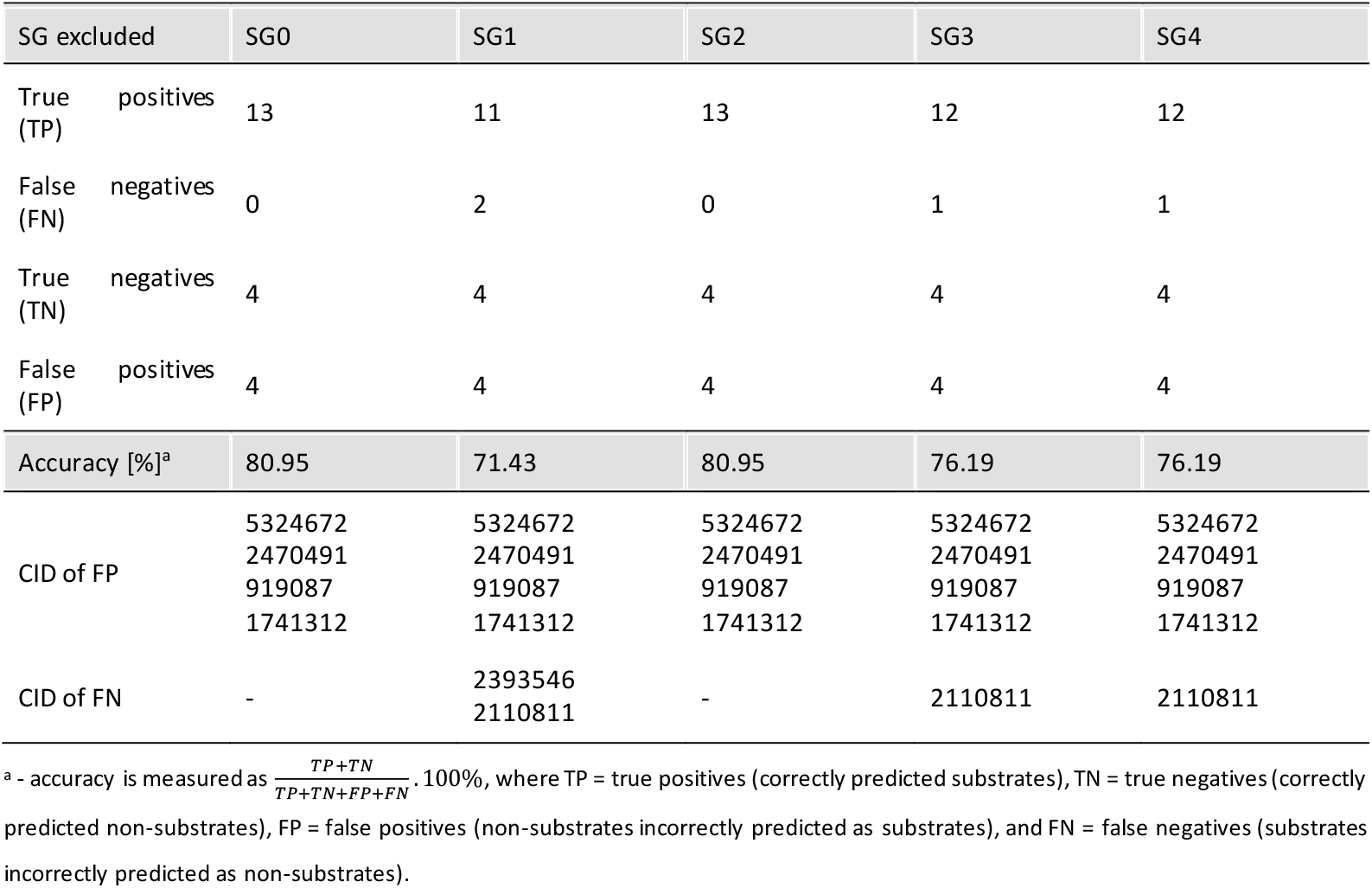
Substrate prediction performance and robustness for HLD family.

Next, we quantified the predictive performance of the derived filters using our target dehalogenase 3U1T (not present among structures used for training), against the testing dataset consisting of 13 experimentally verified substrates and 8 non-substrates (inhibitors), which all can bind the active site and contain appropriate halogenated carbons typical for usual substrates (**Supporting information table 5**). The prediction accuracy ranged from 71-81% depending on the SG excluded from the training, with the average accuracy of 77% (**Table 1**). Analysis of prediction errors revealed a consistent pattern across all substrate group exclusions, with false positives (FP) being the primary source of classification errors. The false positive rate was uniformly 50% (4 out of 8 non-substrates incorrectly predicted as substrates) across all five SG-exclusion experiments, while the false negative (FN) rate averaged only 6.2% (ranging from 0-15.4% depending on the excluded SG). This represents a substantial improvement over the previous work by Daniel et al.,^[33]^ where all inhibitors tested in our study were incorrectly predicted as substrates using the mechanism-based geometric criteria, yielding only 50% overall accuracy. Notably, our method achieves high sensitivity (93.8% average true positive (TP) rate) in identifying actual substrates, with only one substrate (CID 2110811) being consistently misclassified in three out of five SG-exclusion experiments, representing a suitable trade-off for initial screening by proposing some FP candidates for testing rather than missing potential substrates (FN). While reviewing the binding modes of the FN cases, we noted that the reason for their prediction as non-substrates was slightly insufficient accessibility of one pair of atoms in all analyzed conformations. The four consistently misclassified FP non-substrates (CIDs: 5324672, 2470491, 919087, and 1741312) share a common structural feature with competing functional groups (hydroxyl, ketone, amide, or thiazole) being positioned near the catalytic aspartate that allowed simultaneous satisfaction of distance constraints (nucleophilic attack/halogen stabilization) and accessibility thresholds (**Supporting Information Table 8**). Unlike TN, which all contain bulky heterocycles (thiazole, thiadiazole, or benzothiophene) preventing suitable positioning of the leaving groups, the FP ‘substrate mimics’ satisfy the geometric constraints but likely fail to undergo catalysis due to electronic effects. These patterns suggests that while geometric descriptors effectively capture the spatial requirements for catalysis, incorporating electronic parameters could further improve specificity without sacrificing the method’s high sensitivity.

Having established the effectiveness of our approach for the HLD family, we next performed a proof-of-concept transfer to a mechanistically distinct enzyme class (AldR). The AldR family dataset encompassed 9 enzyme structures, including both human and rodent isoforms, with 61 experimentally validated substrates sourced from BRENDA and recent literature (**Supporting Information Table 2**). The substrates were divided into five SG, mainly reflecting their aldehyde or ketone functionalities and carbon chain characteristics, with 36 substrates used for training and 25 for testing. The molecular docking procedure generated between 2,600-9,500 ES-poses only compared to HLD. However, these poses carried larger diversity as illustrated by their clustering into 14-16 distinct binding modes based on their interaction distance profiles. Analogously, the complexity of this dataset is reflected in the generation of 5-8 filters per SG exclusion, demonstrating greater mechanistic diversity compared to the HLD family (**Supporting Information Table 7**). Hence, the best filter was selected based on the number of ES-complexes used for its generation. The geometric filters derived for the AldR family showed remarkable consistency across SG-exclusion experiments, with all filters incorporating the same three key interaction types despite the chemical diversity of substrates. The filters exclusively utilized interactions with the catalytic histidine(HIS-NE2 and HIS-HE2), the catalytic tyrosine (TYR-HH), and the NADPH cofactor (NPH-C4N atom), reflecting the conserved catalytic mechanism across this enzyme family (**Supporting Information Table 9**). The histidine interactions were the most variable among the filters, involving either the imidazole nitrogen (NE2) or its proton (HE2) or both, maintained distances ≤4.3 Å with the substrate’s reactive carbon (C1) and ≤3.2 Å carbonyl oxygen (O1), consistent with its role in proton transfer during catalysis. The tyrosine hydroxyl group (TYR-HH) interactions with the substrate oxygen showed distances ≤4.9 Å, supporting its function in stabilizing the alkoxide intermediate. Notably, the NADPH cofactor interactions (NPH-C4N) with the substrate oxygen maintained tight distance constraints ≤ 3.5 Å (except for filters derived with SG2 exclusion), key to the hydride transfer mechanism. Importantly, the accessibility requirements varied markedly across different filters, ranging from 0.510-1.0 to 0.983-1.0, with most filters requiring minimum accessibility values between 0.68-0.88 for productive binding. This broader accessibility range compared to HLD(≥0.78) suggests that the AldR active sites included in the filter derivation are structurally more heterogeneous, which is supported by the markedly larger number of binding mode clusters per SG-exclusion observed in this family than for HLDs (**Supporting Information Table 7**). However, the consistent involvement of only HIS, TYR, and NADPH atoms demonstrates that substrate diversity is accommodated through subtle variations in positioning and active site perturbations. Overall, both filter components, accessibility and distance, were also rather permissible in contrast to HLD family.

The performance evaluation for the AldR family using an independent test set of 25 verified substrates with target enzyme 3O3R revealed moderate but consistent substrate recognition capabilities across different training set compositions. While the absence of available testing of non-substrates prevented assessment of specificity and FP rates to fully understand method’s performance for this enzyme family, this analysis still presents valuable proof-of-concept for method transferability across different enzyme families. The filters achieved recall rates ranging from 52% to 60% across the five SG-exclusions (**Table 2**). While these recall values are lower than those observed for HLDs (≥85%), they reflect the greater structural diversity of AldR substrates, which include aromatic and aliphatic aldehydes, ketones, and various cyclic compounds spanning different size ranges and electronic properties. The most notable performance variation occurred when SG2 (mostly composed of aromatic aldehydes) was excluded from training, resulting in the lowest recall of 52%, i.e., only 13 of 25 substrates identified. Analysis of the missed substrate candidates from the testing set (CIDs: 7054, 650, and 880) reveals these are aromatic or conjugated aldehydes. Conversely, exclusion of SG0, SG1, and SG3 maintained 60% recall, identifying 15 of 25 test substrates consistently. The ten substrates consistently missed across most experiments (CIDs: 5280598, 5283335, 5283344, 6763, 31289, 8175, 6184, 541, 8494, and 7487) fell into distinct problematic categories: long aliphatic chains (C6-C15) that likely exceed active site dimensions captured by our filters, and ketones that present different steric environments to examples predominantly in the training set. These systematic failures indicate that while our geometric descriptors effectively capture requirements for typical AldR substrates, cases involving extreme size or different carbonyl geometries would require specialized treatment or relaxed thresholds. The generation of larger number of binding mode clusters per SG-exclusion (**Supporting Information Table 7**) compared to HLDs suggests greater mechanistic heterogeneity in substrate recognition. Despite this diversity, the relatively consistent recall rates (52-60%) indicate that our geometric descriptors capture fundamental requirements for AldR substrate binding, though with lower sensitivity than achieved for the more mechanistically homogeneous HLD family. Intrigued by these results, a structural analysis of the docked conformations employed in filter generation was performed. In most cases, the hydroxyl group of the TYR residue of the active site was oriented opposite to the active site cavity, making it less accessible to the substrates (**Figure 3a**). This hydrogen is necessary for the reaction mechanism, since it is this TYR the hydrogen donor for the aldehyde or ketone group of the substrate that is going to be reduced to alcohol. These conformations contributed to the higher permissivity of distances and accessibilities present in the filters generated. In the target enzyme, this hydrogen was oriented towards the opposite side, making it less accessible to the candidates tested (**Figure 3b**). In this family, accessibility seems to be the marked contributing factor for the lower recall present.

**Figure 3.**
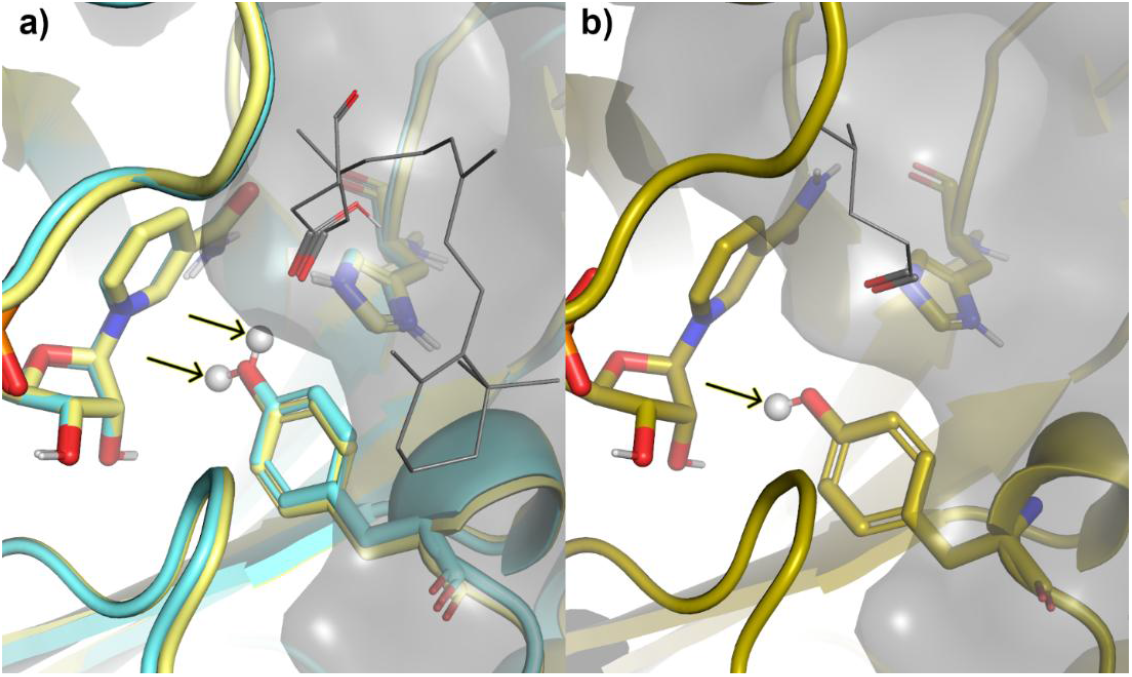
ES-complexes of AldRs. a) Example ES-complexes employed to generate filters in AldRs. Proteins 1ZUA and 1US0 in cartoon representation with the active site residues TYR-49, HIS-111 and cofactor NPH-317 as sticks. Two substrates (CIDs 1112 and 6436082) are represented as lines with the reactive fragments shown as sticks. The hydrogen of TYR-49 (shown as sphere and marked with a black arrow) is pointing in opposite directions in these two enzymes. b) Target protein 3O3R presenting the hydrogen of TYR-49 pointing opposite the cavity, making it less accessible to the candidate substrate (CID 129). Protein represented as cartoon and surface with the active site residues TYR-49, HIS-111 and cofactor NPH-317 as sticks.

**Table 2.** Substrate prediction performance and robustness for AldR family.

| SG excluded | SG0 | SG1 | SG2 | SG3 | SG4 |
| --- | --- | --- | --- | --- | --- |
| Total substrates (TC) | 25 | 25 | 25 | 25 | 25 |
| Identified substrates (IS) | 15 | 15 | 13 | 15 | 14 |
| Recall [%]<sup>a</sup> | 60.00 | 60.00 | 52.00 | 60.00 | 56.00 |
| CID of unidentified substrates | 5280598<br>5283335<br>5283344<br>6763<br>31289<br>8175<br>6184<br>541<br>8494<br>7487 | 5280598<br>5283335<br>5283344<br>6763<br>31289<br>8175<br>6184<br>541<br>8494<br>7487 | 7054<br>5283335<br>5283344<br>6763<br>31289<br>8175<br>6184<br>541<br>8494<br>7487<br>650<br>880 | 5280598<br>5283335<br>5283344<br>6763<br>31289<br>8175<br>6184<br>541<br>8494<br>7487 | 5280598<br>5283335<br>5283344<br>6763<br>31289<br>8175<br>6184<br>541<br>8494<br>7487<br>79014 |

<sup>a</sup>- Recall is measured as $\frac{IS}{TC} \cdot 100\%$ , representing the proportion of actual substrates correctly identified by the geometric filters.

Analysis of false negative predictions across both enzyme families revealed accessibility as the primary limiting factor for accurate substrate identification. In both families, substrates incorrectly classified as non-substrates consistently exhibited accessibility values at the boundary of the defined thresholds. When we implemented a tolerance margin of 0.1 for the accessibility parameter during post-hoc analysis, FN rates decreased, but the FP rates increased (data not shown). This improvement suggests that accessibility calculations, which depend on accurate side chain positioning and conformational sampling during docking, represent the most sensitive component of our geometric filters to differentiate between true substrates and inhibitors. The finding also indicates that incorporating conformational flexibility or ensemble docking approaches could improve the method’s sensitivity. Furthermore, a minimal knowledge about the reaction mechanism is required for the protocol to function successfully, since when all the residues that are described as catalytic in M-CSA were included in the filter selection, the efficiency of the method is dramatically reduced, and often no suitable filters were obtained because the residues that do not interact directly with the substrate were usually further away from the substrate binding site surface and shielded by other atoms. However, when accurate information about the key residues is available, the method showcases its benefits; high accuracy, low computational demands with short running time (the whole protocol can be run in less than a day with a modern laptop), a simple approach, and the results that can be easily interpreted by non-experts.

To contextualize our geometry-based approach within the current landscape of computational substrate prediction, we compared our results with state-of-the-art machine learning methods, which have a very attractive characteristic that most often they do not require the 3D structures of enzymes and substrates, and neither knowledge of the reaction mechanism. Using the ESP and FusionESP web servers, we evaluated predictions for our test datasets, emulating realistic zero-shot use cases where practitioners query public servers with default settings for enzyme families of their interest (**Supporting Information Table 10**). Surprisingly, both models showed severe limitations on our specific test sets. According to the ESP model, all the candidates tested were classified as non-substrates for the enzymes analyzed, yielding a 38.1% accuracy for HLDs, correctly predicting all 8 non-substrates of HDLs (100% specificity) but failing to identify any of the 13 known substrates (0% sensitivity), and 0% recall for AldRs. Similarly, FusionESP model recognized all candidates as non-substrates with high confidence scores. Curiously, both methods exhibited over 90% accuracies on their test sets, implying that they lack either sufficient HLDs and AldRs representation in their training data and/or employ decision thresholds optimized for other enzyme families. For the ESP model, this is in line with the previous observation of significantly reduced accuracy for small molecules that are not present in the training dataset.^[19]^ In case of FusionESP, the model was trained on significant amount of phylogenetic information that could favor the natural well-defined substrates for recognition and conversion of which enzymes have evolved.^[18]^ This could explain the failure in identifying diverse substrates of enzymes with broader specificity, like HLDs and AldRs, which constitutes considerable drawback of such methods for biotechnology-oriented applications where it is desirable to know what new substrates could be converted by a given enzyme. Our geometry-based approach, achieving 71-81% accuracy for HLDs and 52-60% recall for AldRs, demonstrates that mechanistic insights can complement or even surpass purely data-driven methods when dealing with specific enzyme families. Underscoring that even sophisticated ML architecture alone cannot overcome data limitations despite being trained on thousands of examples, where as geometric descriptors grounded in catalytic mechanism and using as few as tens of ES-complexes for training could provide robust predictions even for enzyme families with limited representation in training databases. Additional advantage of our approach lies in its interpretability. Each geometric descriptor directly corresponds to a physical interaction required for catalysis, enabling researchers to understand why a molecule is predicted as a substrate or non-substrate. This interpretability is particularly valuable for enzyme engineering applications, where understanding the structural basis of substrate recognition could be helpful in guiding rational design efforts. Finally, our approach requires only structural information and can be applied to newly discovered enzymes without extensive training data, addressing a critical limitation of data-hungry ML approaches in scenarios with limited experimental validation.

### Conclusion

As enzyme discovery accelerates through metagenomics and protein engineering, the ability to rapidly predict substrates of these enzymes has become a critical bottleneck. Current approaches either require extensive experimental data that is unavailable for newly discovered enzymes, or demand computational resources and expertise that limit their accessibility. Here, we have developed an approach that addresses both limitations simultaneously by automatically deriving interpretable geometric descriptors from minimal training data. The developed and validated a methodology for automatic derivation of geometry-based descriptors from enzyme-substrate complex structures enables substrate prediction without relying on extensive quantum mechanical calculations or requiring large training datasets. Our approach demonstrated robust performance across two mechanistically distinct enzyme families: achieving 77% average accuracy with 94% sensitivity for haloalkane dehalogenases (9 enzymes, 53 substrates) and 57% average recall for aldehyde reductases (9 enzymes, 36 substrates). Notably, these results surpassed state-of-the-art machine learning methods, ESP and FusionESP, which when applied under default web-server thresholds and zero-shot condition to our test sets, both failed to identify any known substrate.

The key advantages of our approach are threefold: i) the geometric descriptors are directly interpretable, corresponding to specific catalytic requirements such as nucleophilic attack distances and halide stabilization interactions; ii) the methodology requires minimal training data—tens of enzyme-substrate complexes rather than thousands—making it applicable to enzyme families with limited experimental characterization; and iii) the computational demands are modest, with the complete protocol executable on a standard laptop within a day. However, several limitations warrant consideration. The method requires 3D structures of enzymes and substrates, knowledge of residues directly interacting with substrates and inherits the accuracy limitations of molecular docking. Also, the performance of the method depends on the consistency of used protein structures, as observed with differential positioning of interacting hydrogen of TYR-49 in aldehyde reductases. Additionally, our analysis identified accessibility calculations as the most sensitive component, suggesting that incorporating conformational flexibility could improve sensitivity.

Future directions include integration of electronic parameters to reduce false positive rates, ensemble docking approaches to better capture conformational diversity, and extension to additional enzyme families. The methodology and datasets are publicly available via Zenodo repository, facilitating application to newly characterized enzymes and supporting enzyme engineering efforts where mechanistic understanding of substrate recognition is essential.

## Acknowledgements

This work was supported by the National Science Centre, Poland (Grant Number 2020/37/N/NZ2/00967). The computations were performed at the Poznan Supercomputing and Networking Center.

