## Supporting information for "Towards automatic derivation of geometry-based descriptors as surrogates for complex computational approaches in enzyme-substrate prediction"

[b] P. Škoda
Department of Software Engineering, Faculty of Mathematics and Physics
Charles University
Malostranské nám. 2/25, 121 16, Praha 1, (Czech Republic)

**Supporting information table 1.** List of substrates from haloalkane dehalogenase family used in this study. For each substrate, the CID ID, name, substrate group (SG) at which it was assigned by similarity and the set at which it was employed. ES-complexes information was gathered from public databases and literature.[33]

| Substrate CID | IUPAC name | SG | Set |
|---|---|---|---|
| 441094 | (3R,6R)-1,3,4,6-tetrachlorocyclohexa-1,4-diene | 0 | Training |
| 7280 | 1,2-dibromo-3-chloropropane | 0 | Training |
| 8101 | 1-bromohexane | 0 | Training |
| 137636 | bromomethylcyclohexane | 0 | Training |
| 8140 | 1-bromooctane | 0 | Training |
| 727 | 1,2,3,4,5,6-hexachlorocyclohexane | 0 | Training |
| 7960 | bromocyclohexane | 0 | Training |
| 5324672 | (1S)-2-chloro-1-(2,4-dichlorophenyl)ethanol | 0 | Test |
| 2392384 | (7R)-2-(chloromethyl)-7-methyl-5,6,7,8-tetrahydro-3H-[1]benzothiolo[2,3-d]pyrimidin-4-one | 0 | Test |
| 2470491 | (E)-2-(1,3-benzothiazol-2-yl)-4-chloro-3-methylbut-2-enenitrile | 0 | Test |
| 713079 | 1-(iodomethyl)-[1,3]thiazolo[3,2-a]benzimidazole | 0 | Test |
| 1666398 | 2-(3-chloropropanoylamino)-4,5,6,7-tetrahydro-1-benzothiophene-3-carboxamide | 0 | Test |
| 691093 | 2-[[3-(chloromethyl)-2,4,6-trimethylphenyl]methylidene]propanedinitrile | 0 | Test |
| 919087 | 2-chloro-1-(1,2,3,4-tetrahydrocarbazol-9-yl)ethanone | 0 | Test |
| 2110811 | 2-chloro-N-(1-cyanocyclohexyl)-N-methylacetamide | 0 | Test |
| 16813824 | 2-chloro-N-(5-phenyl-1,3,4-thiadiazol-2-yl)propanamide | 0 | Test |
| 1484767 | 3-(chloromethyl)-5-(2-chlorophenyl)-1,2-oxazole | 0 | Test |
| 1714004 | 3-chloro-N-(3-cyano-4,5,6,7-tetrahydro-1-benzothiophen-2-yl)propanamide | 0 | Test |
| 1741312 | 3-chloro-N-(3-cyano-4,5-dimethylthiophen-2-yl)propanamide | 0 | Test |
| 943534 | 4-(chloromethyl)-5,7,8-trimethylchromen-2-one | 0 | Test |
| 15758400 | 4-(chloromethyl)-5-hydroxy-7-methylchromen-2-one | 0 | Test |
| 21412 | 4-bromobutyronitrile | 0 | Test |
| 2393546 | 6,8-dibromo-2-(chloromethyl)-1H-quinazolin-4-one | 0 | Test |
| 222275 | 6-(chloromethyl)phenanthridine | 0 | Test |
| 445768 | (R)-1,2-dichloropropane | 1 | Training |
| 17753912 | (S)-1,2-dichloropropane | 1 | Training |
| 7285 | 1,2,3-trichloropropane | 1 | Training |
| 11 | 1,2-dichloroethane | 1 | Training |
| 6564 | 1,2-dichloropropane | 1 | Training |
| 8881 | 1,3-dichloropropane | 1 | Training |
| 8059 | 1,4-dichlorobutane | 1 | Training |
| 7849 | 1-bromo-2-chloroethane | 1 | Training |
| 8006 | 1-bromo-3-chloropropane | 1 | Training |
| 8005 | 1-chlorobutane | 1 | Training |
| 10899 | 1-chloropropane | 1 | Training |
| 6565 | 2,3-dichloroprop-1-ene | 1 | Training |
| 18175 | 2-bromo-1-chloropropane | 1 | Training |
| 6563 | 2-chlorobutane | 1 | Training |
| 34 | 2-chloroethanol | 1 | Training |
| 6361 | 2-chloropropane | 1 | Training |
| 11241 | 3-chloro-2-methylprop-1-ene | 1 | Training |
| 13569 | 4-chlorobutan-1-ol | 1 | Training |
| 8115 | 1-chloro-2-(2-chloroethoxy)ethane | 1 | Training |
| 12353 | 1,5-dichloropentane | 2 | Training |
| 16551 | 1,6-dichlorohexane | 2 | Training |
| 13191 | 1,9-dichlorononane | 2 | Training |
| 13848 | 1-chlorodecane | 2 | Training |
| 12371 | 1-chloroheptane | 2 | Training |
| 10992 | 1-chlorohexane | 2 | Training |

| | | | |
|---|---|---|---|
| 17185 | 1-chlorononane | 2 | Training |
| 8142 | 1-chlorooctane | 2 | Training |
| 10977 | 1-chloropentane | 2 | Training |
| 12347 | 2-chlorooctane | 2 | Training |
| 33016 | 3-chlorohexane | 2 | Training |
| 74828 | 6-chlorohexan-1-ol | 2 | Training |
| 10952 | chlorocyclohexane | 2 | Training |
| 70252 | chlorocyclopentane | 2 | Training |
| 12228454 | (R)-1,2-dibromopropane | 3 | Training |
| 642201 | (S)-1,2-dibromopropane | 3 | Training |
| 7279 | 1,2,3-tribromopropane | 3 | Training |
| 6553 | 1,2-dibromopropane | 3 | Training |
| 8001 | 1,3-dibromopropane | 3 | Training |
| 6555 | 1-bromo-2-methylpropane | 3 | Training |
| 6332 | 1-bromoethane | 3 | Training |
| 7840 | 1-bromopropane | 3 | Training |
| 10898 | 2-bromoethanol | 3 | Training |
| 12308 | 3-bromo-1-propan-1-ol | 3 | Training |
| 7839 | 1,2-dibromoethane | 3 | Test |
| 8002 | 1-bromobutane | 3 | Test |
| 12314 | 1,3-diiodopropane | 4 | Training |
| 12527 | 1-iodohexane | 4 | Training |
| 10559 | 2-iodobutane | 4 | Training |
| 10962 | 1-iodobutane | 4 | Test |
| 33643 | 1-iodopropane | 4 | Test |

**Supporting information table 2.** List of substrates from aldehyde reductase family used in this study. For each substrate, the CID ID, name, substrate group (SG) at which it was assigned by similarity and the set at which it was employed. ES-complexes information was gathered from public databases and literature.[58-72]

| Substrate CID | IUPAC name | SG | Set |
|---|---|---|---|
| 751 | 2,3-dihydroxypropanal | 0 | Training |
| 447466 | (E)-but-2-enal | 0 | Training |
| 79014 | (2R)-2,3-dihydroxypropanal | 0 | Test |
| 7847 | prop-2-enal | 0 | Test |
| 650 | butane-2,3-dione | 0 | Test |
| 7860 | oxaldehyde | 0 | Test |
| 880 | 2-oxopropanal | 0 | Test |
| 6436079 | (2Z,4E,6E,8E)-3,7-dimethyl-9-(2,6,6-trimethylcyclohexen-1-yl)nona-2,4,6,8-tetraenal | 1 | Training |
| 6436082 | (2E,4E,6Z,8E)-3,7-dimethyl-9-(2,6,6-trimethylcyclohexen-1-yl)nona-2,4,6,8-tetraenal | 1 | Training |
| 638015 | (2E,4E,6E,8E)-3,7-dimethyl-9-(2,6,6-trimethylcyclohexen-1-yl)nona-2,4,6,8-tetraenal | 1 | Training |
| 637564 | (2E,4E)-hexa-2,4-dienal | 1 | Training |
| 5283344 | (E)-4-hydroxynon-2-enal | 1 | Test |
| 6445537 | (E)-4-oxonon-2-enal | 1 | Test |
| 5280598 | (2E,6E)-3,7,11-trimethyldodeca-2,6,10-trienal | 1 | Test |
| 5281168 | (E)-hex-2-enal | 1 | Test |
| 5283335 | (E)-non-2-enal | 1 | Test |
| 10667 | naphthalene-1,2-dione | 2 | Training |
| 246876 | (8R,9S,13S,14S)-3-hydroxy-13-methyl-6,7,8,9,11,12,14,15-octahydrocyclopenta[a]phenanthrene-16,17-dione | 2 | Training |
| 6238 | (8R,9S,10R,13S,14S,17R)-17-acetyl-17-hydroxy-10,13-dimethyl-2,6,7,8,9,11,12,14,15,16-decahydro-1H-cyclopenta[a]phenanthren-3-one | 2 | Training |
| 68474 | 2-acetylbenzoic acid | 2 | Training |
| 6813 | 2-benzoylbenzoic acid | 2 | Training |
| 8406 | 2-formylbenzoic acid | 2 | Training |
| 6996 | 2-chlorobenzaldehyde | 2 | Training |
| 46533 | 2-(2-formylphenoxy)acetic acid | 2 | Training |
| 8658 | 2-methoxybenzaldehyde | 2 | Training |
| 12077 | 3-formylbenzoic acid | 2 | Training |
| 11477 | 3-chlorobenzaldehyde | 2 | Training |
| 12088 | 4-formylbenzoic acid | 2 | Training |
| 7726 | 4-chlorobenzaldehyde | 2 | Training |
| 74083 | 9-oxofluorene-1-carboxylic acid | 2 | Training |
| 94715 | (2S,3S,4S,5R)-3,4,5,6-tetrahydroxyoxane-2-carboxylic acid | 2 | Training |
| 219402 | 2-(3,4-dihydroxy-5-oxooxolan-2-yl)-2-hydroxyacetaldehyde | 2 | Training |
| 31261 | pentane-2,4-dione | 2 | Training |
| 8651 | 1,2-diphenylethane-1,2-dione | 2 | Training |
| 30323 | (7S,9S)-9-acetyl-7-[(2R,4S,5S,6S)-4-amino-5-hydroxy-6-methyloxan-2-yl]oxy-6,9,11-trihydroxy-4-methoxy-8,10-dihydro-7H-tetracene-5,12-dione | 2 | Training |
| 31703 | (7S,9S)-7-[(2R,4S,5S,6S)-4-amino-5-hydroxy-6-methyloxan-2-yl]oxy-6,9,11-trihydroxy-9-(2-hydroxyacetyl)-4-methoxy-8,10-dihydro-7H-tetracene-5,12-dione | 2 | Training |
| 3278 | 2-[2,3-dichloro-4-(2-methylidenebutanoyl)phenoxy]acetic acid | 2 | Training |
| 14090 | 2-oxo-2-phenylacetaldehyde | 2 | Training |
| 331384 | 2-[(8S,9S,10R,13S,14S,17S)-10,13-dimethyl-3-oxo-1,2,6,7,8,9,11,12,14,15,16,17-dodecahydrocyclopenta[a]phenanthren-17-yl]-2-oxoacetaldehyde | 2 | Training |
| 14273 | pyridine-2-carbaldehyde | 2 | Training |
| 1112 | 4-oxobutanoic acid | 2 | Training |
| 114839 | (4S,5R)-4,5,6-trihydroxy-2-oxohexanal | 2 | Test |
| 237332 | 5-(hydroxymethyl)furan-2-carbaldehyde | 2 | Test |
| 6763 | phenanthrene-9,10-dione | 2 | Test |
| 240 | benzaldehyde | 2 | Test |

| | | | |
|---|---|---|---|
| 7054 | 1H-indole-2,3-dione | 2 | Test |
| 10371 | pyridine-3-carbaldehyde | 2 | Test |
| 13389 | pyridine-4-carbaldehyde | 2 | Test |
| 13006 | cyclohexane-1,2-dione | 3 | Training |
| 19707 | hexane-2,3-dione | 3 | Training |
| 8175 | 1-decanal | 3 | Test |
| 6184 | 1-hexanal | 3 | Test |
| 31289 | 1-nonanal | 3 | Test |
| 60983 | heptane-2,3-dione | 3 | Test |
| 129 | 4-methylpentanal | 3 | Test |
| 11346 | 1-(2-nitrophenyl)ethenone | 4 | Training |
| 11101 | 2-nitrobenzaldehyde | 4 | Training |
| 7449 | 3-nitrobenzaldehyde | 4 | Training |
| 8494 | 1-(3-nitrophenyl)ethanone | 4 | Test |
| 7487 | 1-(4-nitrophenyl)ethanone | 4 | Test |
| 541 | 4-nitrobenzaldehyde | 4 | Test |

**Supporting information table 3.** List of experimentally verified substrates of haloalkane dehalogenases from literature for each enzyme used in the training set. The corresponding substrates of each enzyme are marked with an X.

| Enzymes PDB IDs | | | | | | | | | |
|---|---|---|---|---|---|---|---|---|---|
| Substrate CID | 1MJ5 | 1CQW | 3A2M | 2QVB | 4K2A | 4E46 | 4HZG | 4C6H | 4H77 |
| 11 | | X | X | | | | | | |
| 34 | X | | | | | | | | |
| 727 | | | | | | | | | X |
| 6332 | X | | | | | | | | |
| 6553 | | | | | | X | | | |
| 6555 | X | | | | | | | | |
| 6361 | | | | | | | | X | |
| 6563 | X | | | | | | | X | |
| 6564 | | | | | | X | | | |
| 6565 | X | X | X | X | X | | | | |
| 7279 | X | X | X | X | X | X | | | |
| 7280 | | X | | | X | | | | |
| 7285 | | X | X | | | X | X | | |
| 7839 | X | X | X | X | X | X | | X | |
| 7840 | X | | | | | X | | | |
| 7849 | X | X | X | X | X | | | | |
| 7960 | X | X | X | X | X | | | X | |
| 8001 | X | X | X | X | X | X | | X | |
| 8002 | X | X | X | X | X | | | | |
| 8005 | X | X | X | X | X | X | | | |
| 8006 | X | X | X | X | X | | | | |
| 8059 | X | | | | | | | X | |
| 8101 | X | X | X | X | X | | | X | |
| 8115 | X | X | X | X | X | | | | |
| 8140 | | | | | | | | X | |
| 8142 | X | | | | | | | | |
| 8881 | X | X | X | X | | X | | | |
| 10559 | X | X | X | X | X | | | | |
| 10898 | | | | | | X | | | |
| 10899 | X | | | | | X | | | |
| 10952 | X | X | X | | | | | | |
| 10962 | X | X | X | X | X | X | | | |
| 10977 | X | | | | | | | | |
| 10992 | X | X | X | X | X | | | | |
| 11241 | X | X | X | X | X | | | | |
| 12308 | | | | | | X | | | |
| 12314 | X | X | X | | X | | | | |
| 12347 | X | | | | | | | | |
| 12353 | X | X | X | X | X | | | | |
| 12371 | X | | | | | | | | |
| 12527 | X | X | X | X | X | | | | |
| 13191 | X | | | | | | | | |
| 13569 | X | | | | | | | | |
| 13848 | X | | | | | | | | |
| 16551 | X | | | | | | | X | |
| 17185 | X | | | | | | | | |
| 18175 | X | X | X | X | X | | | | |
| 21412 | X | X | X | X | X | | | | |

| | | | | | |
|---|---|---|---|---|---|
| 33016 | X | | | | |
| 33643 | X | X | X | X | X |
| 70252 | X | X | X | X | X |
| 74828 | X | | | | |
| 137636 | X | X | | | X |
| 441094 | X | | | | |
| 445768 | | | X | | |
| 642201 | X | X | X | X | X |
| 12228454 | X | X | X | X | X |
| 17753912 | | | X | | |

**Supporting information table 4.** List of experimentally verified substrates of aldehyde reductase from literature for each enzyme used in the training set. The corresponding substrates of each enzyme are marked with an X.

| Enzymes PDB/UNIPROT IDs | | | | | | | | | |
|---|---|---|---|---|---|---|---|---|---|
| Substrate CID | 3FX4 | 2BP1 | 1ZUA | 1US0 | 2C91 | P14550 | 4GAC | 1FRB | 1C9W |
| 129 | | | X | | | | | | |
| 240 | | | | X | | | | | X |
| 541 | | | | X | | X | | | X |
| 751 | X | | X | X | | X | X | X | |
| 880 | | | | X | | X | X | | X |
| 1112 | | X | | X | X | X | | | |
| 3278 | | | | | | X | | | |
| 6238 | | | | | | | | | X |
| 6763 | | X | | X | | X | | | |
| 6813 | | X | | | | | | | |
| 6996 | | | | X | | X | | | |
| 7054 | | X | | X | | | | | |
| 7449 | | | | X | | X | | | |
| 7487 | | | | X | | X | | | X |
| 7726 | | | | X | | X | | | |
| 7847 | | | X | X | | X | | | |
| 7860 | | | | X | | X | | | |
| 8406 | | X | | X | | | | | |
| 8494 | | | | X | | X | | | |
| 8651 | | | | X | | X | | | |
| 8658 | | X | | | | | | | |
| 10371 | X | | X | X | | X | | | X |
| 10667 | | | | X | | X | | | |
| 11101 | | | | X | | X | | | |
| 11346 | | | | X | | X | | | |
| 11477 | | | | X | | X | | | |
| 12077 | | | | X | | X | | | |
| 12088 | | | | X | | X | | | |
| 13006 | | | | X | | X | | | |
| 13389 | | | | X | | X | | | |
| 14090 | | | | X | | X | | | X |
| 14273 | | | | X | | X | | | |
| 19707 | | | | X | | X | | | |
| 30323 | | | | X | | X | | | X |
| 31261 | | | | | | | | | X |
| 31703 | | | | X | | X | | | |
| 46533 | | X | | | | | | | |
| 68474 | | X | | | | | | | |
| 74083 | | X | | | | | | | |
| 94715 | | | | X | | X | X | | |

| | | | | | | | | | |
|---|---|---|---|---|---|---|---|---|---|
| 114839 | | | | | | | X | | |
| 219402 | | | | | | | X | | |
| 246876 | | X | | X | | X | | | |
| 331384 | | | | | | | | | X |
| 447466 | | | X | | | | | | |
| 637564 | | | X | X | | | | | |
| 638015 | | | X | | | | | | |
| 5281168 | | | X | X | | | | | |
| 5283344 | | | X | X | | X | | | |
| 6436079 | | | X | | | | | | |
| 6436082 | | | X | | | | | | |
| 6445537 | | | X | | | | | | |

**Supporting information table 5.** Target enzymes employed for each family with its corresponding candidate substrates used in the testing set. For Haloalkane dehalogenase family, the candidates are separated in verified substrates (positives) and inhibitors (negatives). All the candidates presented were experimentally verified in previous studies.

| Enzyme family | Target PDB | Candidate CID | Candidate IUPAC name | Substrate group (SG) | Candidate type |
|---|---|---|---|---|---|
| Haloalkane dehalogenase | 3U1T | 5324672 | (1S)-2-chloro-1-(2,4-dichlorophenyl)ethanol | 0 | Negative |
| | | 2392384 | (7R)-2-(chloromethyl)-7-methyl-5,6,7,8-tetrahydro-3H-[1]benzothiolo[2,3-d]pyrimidin-4-one | 0 | Positive |
| | | 2470491 | (E)-2-(1,3-benzothiazol-2-yl)-4-chloro-3-methylbut-2-enenitrile | 0 | Negative |
| | | 713079 | 1-(iodomethyl)-[1,3]thiazolo[3,2-a]benzimidazole | 0 | Negative |
| | | 1666398 | 2-(3-chloropropanoylamino)-4,5,6,7-tetrahydro-1-benzothiophene-3-carboxamide | 0 | Negative |
| | | 691093 | 2-[[3-(chloromethyl)-2,4,6-trimethylphenyl]methylidene]propanedinitrile | 0 | Positive |
| | | 919087 | 2-chloro-1-(1,2,3,4-tetrahydrocarbazol-9-yl)ethanone | 0 | Negative |
| | | 2110811 | 2-chloro-N-(1-cyanocyclohexyl)-N-methylacetamide | 0 | Positive |
| | | 16813824 | 2-chloro-N-(5-phenyl-1,3,4-thiadiazol-2-yl)propanamide | 0 | Negative |
| | | 1484767 | 3-(chloromethyl)-5-(2-chlorophenyl)-1,2-oxazole | 0 | Positive |
| | | 1714004 | 3-chloro-N-(3-cyano-4,5,6,7-tetrahydro-1-benzothiophen-2-yl)propanamide | 0 | Negative |
| | | 1741312 | 3-chloro-N-(3-cyano-4,5-dimethylthiophen-2-yl)propanamide | 0 | Negative |
| | | 943534 | 4-(chloromethyl)-5,7,8-trimethylchromen-2-one | 0 | Positive |
| | | 15758400 | 4-(chloromethyl)-5-hydroxy-7-methylchromen-2-one | 0 | Positive |
| | | 21412 | 4-bromobutyronitrile | 0 | Positive |
| | | 2393546 | 6,8-dibromo-2-(chloromethyl)-1H-quinazolin-4-one | 0 | Positive |
| | | 222275 | 6-(chloromethyl)phenanthridine | 0 | Positive |
| | | 7839 | 1,2-dibromoethane | 3 | Positive |
| | | 8002 | 1-bromobutane | 3 | Positive |
| | | 10962 | 1-iodobutane | 4 | Positive |
| | | 33643 | 1-iodopropane | 4 | Positive |
| Aldehyde reductase | 3O3R | 79014 | (2R)-2,3-dihydroxypropanal | 0 | Positive |
| | | 7847 | prop-2-enal | 0 | Positive |
| | | 650 | butane-2,3-dione | 0 | Positive |
| | | 7860 | oxaldehyde | 0 | Positive |
| | | 880 | 2-oxopropanal | 0 | Positive |
| | | 5283344 | (E)-4-hydroxynon-2-enal | 1 | Positive |
| | | 6445537 | (E)-4-oxonon-2-enal | 1 | Positive |
| | | 5280598 | (2E,6E)-3,7,11-trimethyldodeca-2,6,10-trienal | 1 | Positive |
| | | 5281168 | (E)-hex-2-enal | 1 | Positive |
| | | 5283335 | (E)-non-2-enal | 1 | Positive |
| | | 114839 | (4S,5R)-4,5,6-trihydroxy-2-oxohexanal | 2 | Positive |
| | | 237332 | 5-(hydroxymethyl)furan-2-carbaldehyde | 2 | Positive |
| | | 6763 | phenanthrene-9,10-dione | 2 | Positive |
| | | 240 | benzaldehyde | 2 | Positive |
| | | 7054 | 1H-indole-2,3-dione | 2 | Positive |
| | | 10371 | pyridine-3-carbaldehyde | 2 | Positive |
| | | 13389 | pyridine-4-carbaldehyde | 2 | Positive |
| | | 8175 | 1-decanal | 3 | Positive |
| | | 6184 | 1-hexanal | 3 | Positive |
| | | 31289 | 1-nonanal | 3 | Positive |
| | | 60983 | heptane-2,3-dione | 3 | Positive |
| | | 129 | 4-methylpentanal | 3 | Positive |
| | | 8494 | 1-(3-nitrophenyl)ethanone | 4 | Positive |
| | | 7487 | 1-(4-nitrophenyl)ethanone | 4 | Positive |
| | | 541 | 4-nitrobenzaldehyde | 4 | Positive |

**Supporting information table 6.** List of residues and atoms in each family that are necessary for the catalytic reaction according to the description of their reaction mechanism. For each atom, the maximum distance and minimum accessibility considered are presented. The residue numbers correspond to the numbering ID in the PDB 3U1T for haloalkane dehalogenase and 3O3R for aldehyde reductase.

| Enzyme family | Residue name | Residue number in target PDB | Atom name | Maximum distance [Å] | Minimum accessibility [fraction] |
|---|---|---|---|---|---|
| Haloalkane dehalogenase | ASN | 78 | HD21 | 6 | 0.5 |
| | | | HD22 | 6 | 0.5 |
| | ASP | 144 | OD1 | 6 | 0.5 |
| | | | OD2 | 6 | 0.5 |
| | TRP | 145 | HE1 | 6 | 0.5 |
| Aldehyde reductase | HIS | 111 | ND1 | 5 | 0.5 |
| | | | NE2 | 5 | 0.5 |
| | | | HD1 | 5 | 0.5 |
| | | | HE2 | 5 | 0.5 |
| | TYR | 49 | OH | 6 | 0.5 |
| | | | HH | 6 | 0.5 |
| | NPH | 317 | C4N | 6 | 0.5 |

**Supporting information table 7.** Statistics on the datasets used to derive filters when excluding each substrate group (SG).

| Enzyme family | SG excluded | # of enzymes used | # of substrates used | # of docked poses | # binding mode clusters | # of derived filters | ID of the best filter | # of unique ES-complexes used to derive the best filter |
|---|---|---|---|---|---|---|---|---|
| Haloalkane dehalogenase | 0 | 8 | 46 | 24943 | 2 | 1 | 0 | 65 |
| | 1 | 8 | 34 | 14255 | 3 | 1 | 1 | 44 |
| | 2 | 9 | 39 | 21970 | 3 | 2 | 2 | 63 |
| | 3 | 9 | 43 | 21060 | 4 | 1 | 1 | 58 |
| | 4 | 9 | 50 | 25040 | 5 | 2 | 2 | 65 |
| Aldehyde reductase | 0 | 7 | 34 | 8997 | 14 | 5 | 2 | 21 |
| | 1 | 9 | 32 | 9572 | 15 | 5 | 2 | 22 |
| | 2 | 6 | 11 | 2678 | 14 | 7 | 8 | 8 |
| | 3 | 9 | 34 | 9007 | 15 | 5 | 1 | 22 |
| | 4 | 9 | 33 | 9470 | 16 | 6 | 1 | 21 |

**Supporting information table 8.** Filters generated when excluding each substrate group (SG) for haloalkane dehalogenase family. For each filter, the atom pairs of the ES-complexes obtained are presented with their corresponding intervals. For the case of ASN and ASP, the atoms are interchangeable, hence, the names are unified as HD2 and OD respectively.

| SG excluded | Selected filter ID | Enzyme's atom | Fragment's atom | Measurement | Interval |
|---|---|---|---|---|---|
| 0 | 0 | ASN-HD2 | X1 | distance | 0.0 – 3.721 |
| | | | | accessibility | 0.942 – 1.0 |
| | | ASP-OD | C1 | distance | 0.0 – 3.419 |
| | | | | accessibility | 0.775 – 1.0 |
| | | TRP-HE1 | X1 | distance | 0.0 – 3.164 |
| | | | | accessibility | 0.917 – 1.0 |
| 1 | 1 | ASN-HD2 | X1 | distance | 0.0 – 3.381 |
| | | | | accessibility | 0.958 – 1.0 |
| | | ASP-OD | C1 | distance | 0.0 – 3.455 |
| | | | | accessibility | 0.775 – 1.0 |
| | | TRP-HE1 | X1 | distance | 0.0 – 2.912 |
| | | | | accessibility | 0.925 – 1.0 |
| 2 | 2 | ASN-HD2 | X1 | distance | 0.0 – 3.140 |
| | | | | accessibility | 0.942 – 1.0 |
| | | ASP-OD | C1 | distance | 0.0 – 3.434 |
| | | | | accessibility | 0.817 – 1.0 |
| | | TRP-HE1 | X1 | distance | 0.0 – 2.985 |
| | | | | accessibility | 0.917 – 1.0 |
| 3 | 1 | ASN-HD2 | X1 | distance | 0.0 – 3.312 |
| | | | | accessibility | 0.958 – 1.0 |
| | | ASP-OD | C1 | distance | 0.0 – 3.455 |
| | | | | accessibility | 0.800 – 1.0 |
| | | TRP-HE1 | X1 | distance | 0.0 – 2.997 |
| | | | | accessibility | 0.917 – 1.0 |
| 4 | 2 | ASN-HD2 | X1 | distance | 0.0 – 3.082 |
| | | | | accessibility | 0.958 – 1.0 |
| | | ASP-OD | C1 | distance | 0.0 – 3.455 |
| | | | | accessibility | 0.800 – 1.0 |
| | | TRP-HE1 | X1 | distance | 0.0 – 2.912 |
| | | | | accessibility | 0.917 – 1.0 |

**Supporting information table 9.** Filters generated when excluding each substrate group (SG) for aldehyde reductase family. For each filter, the atom pairs of the ES-complexes obtained are presented with their corresponding intervals.

| SG excluded | Selected filter ID | Enzyme's atom | Fragment's atom | Measurement | Interval |
|---|---|---|---|---|---|
| 0 | 2 | HIS-NE2 | C1 | distance | 0.0 – 3.959 |
| | | | | accessibility | 0.510 – 1.0 |
| | | HIS-HE2 | O1 | distance | 0.0 – 3.030 |
| | | | | accessibility | 0.679 – 1.0 |
| | | TYR-HH | O1 | distance | 0.0 – 4.544 |
| | | | | accessibility | 0.983 – 1.0 |
| | | NPH-C4N | O1 | distance | 0.0 – 3.502 |
| | | | | accessibility | 0.825 – 1.0 |
| 1 | 2 | HIS-NE2 | C1 | distance | 0.0 – 4.254 |
| | | | | accessibility | 0.510 – 1.0 |
| | | HIS-HE2 | O1 | distance | 0.0 – 3.176 |
| | | | | accessibility | 0.764 – 1.0 |
| | | TYR-HH | O1 | distance | 0.0 – 4.508 |
| | | | | accessibility | 0.983 – 1.0 |
| | | NPH-C4N | O1 | distance | 0.0 – 3.502 |
| | | | | accessibility | 0.825 – 1.0 |
| 2 | 8 | HIS-HE2 | O1 | distance | 0.0 – 2.391 |
| | | | | accessibility | 0.983 – 1.0 |
| | | TYR-HH | O1 | distance | 0.0 – 4.015 |
| | | | | accessibility | 0.975 – 1.0 |
| | | NPH-C4N | O1 | distance | 0.0 – 4.120 |
| | | | | accessibility | 0.850 – 1.0 |
| 3 | 1 | HIS-NE2 | C1 | distance | 0.0 – 4.254 |
| | | | | accessibility | 0.510 – 1.0 |
| | | HIS-HE2 | O1 | distance | 0.0 – 3.176 |
| | | | | accessibility | 0.700 – 1.0 |
| | | TYR-HH | O1 | distance | 0.0 – 4.305 |
| | | | | accessibility | 0.983 – 1.0 |
| | | NPH-C4N | O1 | distance | 0.0 – 3.502 |
| | | | | accessibility | 0.825 – 1.0 |
| 4 | 1 | HIS-NE2 | C1 | distance | 0.0 – 3.900 |
| | | | | accessibility | 0.612 – 1.0 |
| | | TYR-HH | O1 | distance | 0.0 – 4.900 |
| | | | | accessibility | 0.883 – 1.0 |
| | | NPH-C4N | O1 | distance | 0.0 – 3.584 |
| | | | | accessibility | 0.825 – 1.0 |

**Supporting information table 10.** Substrate prediction results obtained with ESP and FusionESP models, using their respective web-servers, for the test sets employed in this study.

| PDB id of target enzyme | Substrate CID | ESP model predictions | | FusionESP model predictions | |
|---|---|---|---|---|---|
| | | Is substrate | Score | Is substrate | Confidence score |
| 3U1T | 7839 | No | 0.05 | No | 0.6315 |
| | 8002 | No | 0.03 | No | 0.8867 |
| | 10962 | No | 0.04 | No | 0.8475 |
| | 21412 | No | 0.02 | No | 0.8448 |
| | 33643 | No | 0.06 | No | 0.9786 |
| | 222275 | No | 0.06 | No | 0.8868 |
| | 691093 | No | 0.01 | No | 0.7861 |
| | 713079 | No | 0.06 | No | 0.9995 |
| | 919087 | No | 0.11 | No | 0.8484 |
| | 943534 | No | 0.05 | No | 1.2066 |
| | 1484767 | No | 0.03 | No | 1.0327 |
| | 1666398 | No | 0.07 | No | 0.6199 |
| | 1714004 | No | 0.03 | No | 0.6363 |
| | 1741312 | No | 0.02 | No | 0.6522 |
| | 2110811 | No | 0.01 | No | 0.7839 |
| | 2392384 | No | 0.05 | No | 0.7333 |
| | 2393546 | No | 0.05 | No | 0.9722 |
| | 2470491 | No | 0.02 | No | 0.7443 |
| | 5324672 | No | 0.04 | No | 0.9900 |
| | 15758400 | No | 0.08 | No | 1.1226 |
| | 16813824 | No | 0.05 | No | 0.7394 |
| 3O3R | 129 | No | 0.04 | No | 0.7189 |
| | 240 | No | 0.01 | No | 0.9884 |
| | 541 | No | 0.01 | No | 0.9039 |
| | 650 | No | 0.19 | No | 0.8954 |
| | 880 | No | 0.02 | No | 0.7447 |
| | 6184 | No | 0.01 | No | 0.7050 |
| | 6763 | No | 0.04 | No | 0.9860 |
| | 7054 | Uncertain | 0.52 | No | 0.7626 |
| | 7487 | No | 0.31 | No | 0.7954 |
| | 7847 | No | 0.03 | No | 0.8838 |
| | 7860 | No | 0.02 | No | 1.1260 |
| | 8175 | No | 0.02 | No | 0.8311 |
| | 8494 | No | 0.19 | No | 0.8699 |
| | 10371 | No | 0.01 | No | 0.8144 |
| | 13389 | No | 0.01 | No | 0.8569 |
| | 31289 | No | 0.01 | No | 0.8075 |
| | 60983 | No | 0.02 | No | 0.8155 |
| | 79014 | No | 0.02 | No | 1.0046 |
| | 114839 | No | 0.01 | No | 0.8900 |
| | 237332 | No | 0.01 | No | 0.9805 |
| | 5280598 | No | 0.02 | No | 0.5909 |
| | 5281168 | No | 0.05 | No | 0.7659 |
| | 5283335 | No | 0.37 | No | 0.9138 |
| | 5283344 | No | 0.09 | No | 0.9550 |
| | 6445537 | No | 0.07 | No | 1.0262 |

**Supporting information references:**
